# Excessive *O* - GlcNAcylation causes heart failure and sudden death

**DOI:** 10.1101/2020.02.11.943910

**Authors:** Priya Umapathi, Partha S. Banerjee, Natasha E. Zachara, Neha Abrol, Qinchuan Wang, Olurotimi O. Mesubi, Elizabeth D. Luczak, Yuejin Wu, Jonathan M. Granger, An-Chi Wei, Oscar E. Reyes Gaido, Liliana Florea, C. Conover Talbot, Gerald W. Hart, Mark E. Anderson

**Affiliations:** Division of Cardiology, The Johns Hopkins University School of Medicine, Baltimore, MD 21205, USA; Department of Medicine, The Johns Hopkins University School of Medicine, Baltimore, MD 21205, USA; Department of Biological Chemistry, The Johns Hopkins University School of Medicine, Baltimore, MD 21205, USA; The Complex Carbohydrate Research Center and Department of Biochemistry and Molecular Biology, Univ. of Georgia, Athens GA 30602, USA; Department of Physiology, The Johns Hopkins University School of Medicine, Baltimore, MD 21205, USA; Computational Biology Consulting Core, The Johns Hopkins University School of Medicine, Baltimore, MD 21205, USA; Institute for Basic Biomedical Sciences, The Johns Hopkins University School of Medicine, Baltimore, MD 21205, USA; Department of Electrical Engineering, Graduate Institute of Biomedical and Bioinformatics, National Taiwan University, Taiwan

**Keywords:** Dilated cardiomyopathy, O-GlcNAcylation, mouse model, mitochondrial energetics, pressure overload

## Abstract

**Background:** Heart failure is a leading cause of death worldwide and is associated with the rising prevalence of obesity, hypertension and diabetes. *O-*GlcNAcylation, a post-translational modification of intracellular proteins, serves as a potent transducer of cellular stress. Failing myocardium is marked by increased *O*-GlcNAcylation, but it is unknown if excessive *O-*GlcNAcylation contributes to cardiomyopathy and heart failure. The total levels of *O-*GlcNAcylation are determined by nutrient and metabolic flux, in addition to the net activity of two enzymes, *O-*GlcNAc transferase (OGT) and *O*-GlcNAcase (OGA).

**Methods:** We developed two new transgenic mouse models with myocardial overexpression of OGT and OGA to control O-GlcNAclyation independent of pathological stress.

**Results:** We found that OGT transgenic hearts showed increased *O-*GlcNAcylation, and developed severe dilated cardiomyopathy, ventricular arrhythmias and premature death. In contrast, OGA transgenic hearts had *O-*GlcNAcylation and cardiac function similar to wild type littermate controls. However, OGA trangenic hearts were resistant to pathological stress induced by pressure overload and had attenuated myocardial *O-*GlcNAcylation levels, decreased pathological hypertrophy and improved systolic function. Interbreeding OGT with OGA transgenic mice rescued cardiomyopathy and premature death despite persistant elevation of myocardial OGT. Transcriptomic and functional studies revealed disrupted mitochondrial energetics with impairment of complex I activity in hearts from OGT transgenic mice. Complex I activity was rescued by OGA transgenic interbreeding, suggesting an important role for mitochondrial complex I in *O*-GlcNAc mediated cardiac pathology.

**Conclusions:** Our data provide evidence that excessive *O-*GlcNAcylation causes cardiomyopathy, at least in part, due to defective energetics. Enhanced OGA activity is well tolerated and attenuation of *O-*GlcNAcylation is an effective therapy against pressure overload induced heart failure. Attenuation of excessive *O-*GlcNAcylation may represent a novel therapeutic approach for cardiomyopathy.

**Clinical Perspective:** *What is new?:*

- Cardiomyopathy from diverse causes is marked by increased *O*-GlcNAcylation. Here we provide new genetic mouse models to control myocardial *O*-GlcNAcylation independent of pathological stress.
- Genetically increased myocardial *O-*GlcNAcylation causes progressive dilated cardiomyopathy and premature death, while genetic reduction of myocardial *O*-GlcNAcylation is protective against pathological hypertrophy caused by transaortic banding.
- Excessive myocardial *O*-GlcNAcylation decreases activity and expression of mitochondrial complex I.

*What are the clinical implications?:*

- Increased myocardial *O-*GlcNAcylation has been shown to be associated with a diverse range of clinical heart failure including aortic stenosis, hypertension, ischemia and diabetes.
- Using novel genetic mouse models we have provided new proof of concept data that excessive *O-*GlcNAcylation is sufficient to cause cardiomyopathy.
- We have shown myocardial over-expression of *O*-GlcNAcase, an enzyme that reverses *O*-GlcNAcylation, is well tolerated at baseline, and improves myocardial responses to pathological stress.
- Our findings suggest reversing excessive myocardial *O-*GlcNAcylation could benefit diverse etiologies of heart failure.

## Introduction

Failing myocardium from model animals, and patients is marked by increased protein *O*-GlcNAcylation^1^. However, it is unknown if excessive *O-*GlcNAcylation is a cause or consequence of cardiomyopathy. The hexosamine biosynthesis pathway is a metabolic sensor that utilizes glucose, amino and fatty acids to synthesize UDP-GlcNAc (Uridine Diphosphate N-acetylglucosamine), the substrate for OGT. *O-*GlcNAc is cycled on and off proteins by the activity of two enzymes: OGT (*O-*GlcNAc transferase), which adds GlcNAc from UDP-GlcNAc to proteins, and OGA (*O-*GlcNAcase), which removes UDP-GlcNAc from proteins^2^. On one hand, *O-*GlcNAcylation is a highly conserved stress response, and constitutive OGT knock out is embryonically lethal^3^. Furthermore, inducible loss of myocardial OGT in adult mice causes increased susceptibility to myocardial injury^2^, indicating that *O-*GlcNAcylation is necessary, and suggesting that elevated *O-*GlcNAcylation can be beneficial. On the other hand, excessive *O*-GlcNAcylation is suspected to contribute to myocardial dysfunction in diabetes and hyperglycemia^1, 4^, and OGT inhibitors can reverse or prevent pathological myocardial hypertrophy^1^. Thus, the role of *O-*GlcNAcylation in cardiomyopathy remains a major unresolved question with important implications for the development of new heart failure therapies.

The complexity of myocardial responses to pathological stress, and lack of genetic tools to control *O-*GlcNAcylation levels in vivo, independent of glucose or pathological stress, has limited understanding of the role of increased *O-*GlcNAcylation in cardiomyopathy. We developed novel mouse models to independently control *O-*GlcNAcylation levels in myocardium, and directly test the hypothesis that excessive *O-*GlcNAcylation causes or contributes to cardiomyopathy. Here, we report transgenic myocardial OGT overexpression (OGT TG) causes increased *O-*GlcNAcylation, dilated cardiomyopathy, and premature death. In contrast, transgenic myocardial OGA overexpression (OGA TG) does not cause cardiomyopathy. However, OGA TG mice had reduced myocardial *O-*GlcNAcylation, and were protected against cardiomyopathy due to transaortic banding surgery, a model of acquired pathological myocardial hypertrophy^5, 6^. Interbreeding of OGT TG with OGA TG mice reduced cardiac *O-*GlcNAcylation toward WT levels, rescued dilated cardiomyopathy, and prevented premature death seen in the OGT TG mice. We identified reduced expression of genes and proteins important for oxidative phosphorylation, and impaired energetics in OGT TG hearts; these patterns were restored to near WT levels in hearts from OGT TG x OGA TG interbred mice. Taken together, these data show that excessive *O-*GlcNAcylation is sufficient to cause severe cardiomyopathy, heart failure, and premature mortality. Our findings identify novel targets likely to explain, at least in part, the deleterious effects of excessive myocardial *O-*GlcNAcylation, and suggest reducing myocardial *O-*GlcNAcylation could be a successful therapeutic approach for cardiomyopathy and heart failure.

## Methods

### Animal Models

All the experiments were carried out in accordance with the guidelines of Institutional Animal Care and Use Committee (M017M290). Mice used in these studies were a mixture of male and female animals 7-20 weeks of age unless otherwise noted in the figure legend. We performed our studies on male and female mice with C57BL/6J backgrounds at approximately 3–5 months of age.

#### OGT TG mice

Human cDNA encoding the nucleocytoplasmic variant of the human OGT gene was fused with a C-terminal Myc epitope tag. The resulting construct was cloned into the pBS-αMHC-script-hGH vector for myocardial expression. Pronuclear injections of linearized DNA (digested with NotI) were performed in the Johns Hopkins Transgenic Mouse Core Facility and embryos implanted into pseudo*-*pregnant females to generate C57Bl6/J F1 mice. Insertion of the transgene into the mouse genome was confirmed by PCR analysis (supplement) using the forward primer, 5’-GGA CTT CAC ATA GAA GCC TAG C-3’, and reverse primer, 5’-CAC TGC GAA CAC AGT ACA AAT C--3’, producing a product of 500 base pairs.

#### OGA TG mice

Human cDNA encoding the long form of the Meningioma Expressed Antigen 5 gene (*O*-GlcNAcase, approved symbol OGA) was fused with a N-terminal HA epitope tag. The resulting construct was cloned into the pBS-αMHC-script-hGH vector for myocardial expression. Pronuclear injections of linearized DNA (digested with NotI) were performed in the Johns Hopkins Transgenic Mouse Core Facility and embryos implanted into pseudo*-*pregnant females to generate C57Bl6/J F1 mice. Insertion of the transgene into the mouse genome was confirmed by PCR analysis (supplement) using the forward primer, 5’-TGGTCAGGATCTCTAGATTGGT-3’ and reverse primer, 5’-TCATAAGTTGCTCAGCTTCCTC-3’, producing a product of 850 base pairs. Procedures used for transaortic banding, murine echocardiography, arrhythmia monitoring thru telemetry implant, western blot, OGT and OGA activity assays, mitochondrial isolation, complex I activity assay, Seahorse mitochondrial bioenergetics studies and ventricular myocyte calcium homeostasis followed those previously published and are detailed in the Extended Methods Supplement.

## Results

### Myocardial-targeted OGA overexpression protects against left ventricular hypertrophy and heart failure

Elevated myocardial *O-*GlcNAcylation has been reported in multiple models of cardiomyopathy^1, 7-12^. We first asked if transaortic banding surgery, a validated model of pathologically increased left ventricular afterload^5^, resulted in augmented myocardial *O*-GlcNAcylation (see Methods). We found robust elevation in total *O-*GlcNAcylation in hearts from C57Bl/6J mice with transaortic banding compared to sham operated mice (Fig 1a and b). These results were consistent with previous reports of increased *O*-GlcNAcylation in hearts subjected to pathological stress^1, 12^. Based on these findings, we next asked if attenuation of *O-*GlcNAcylation during sustained cardiac stress could be beneficial.

**Fig 1.**
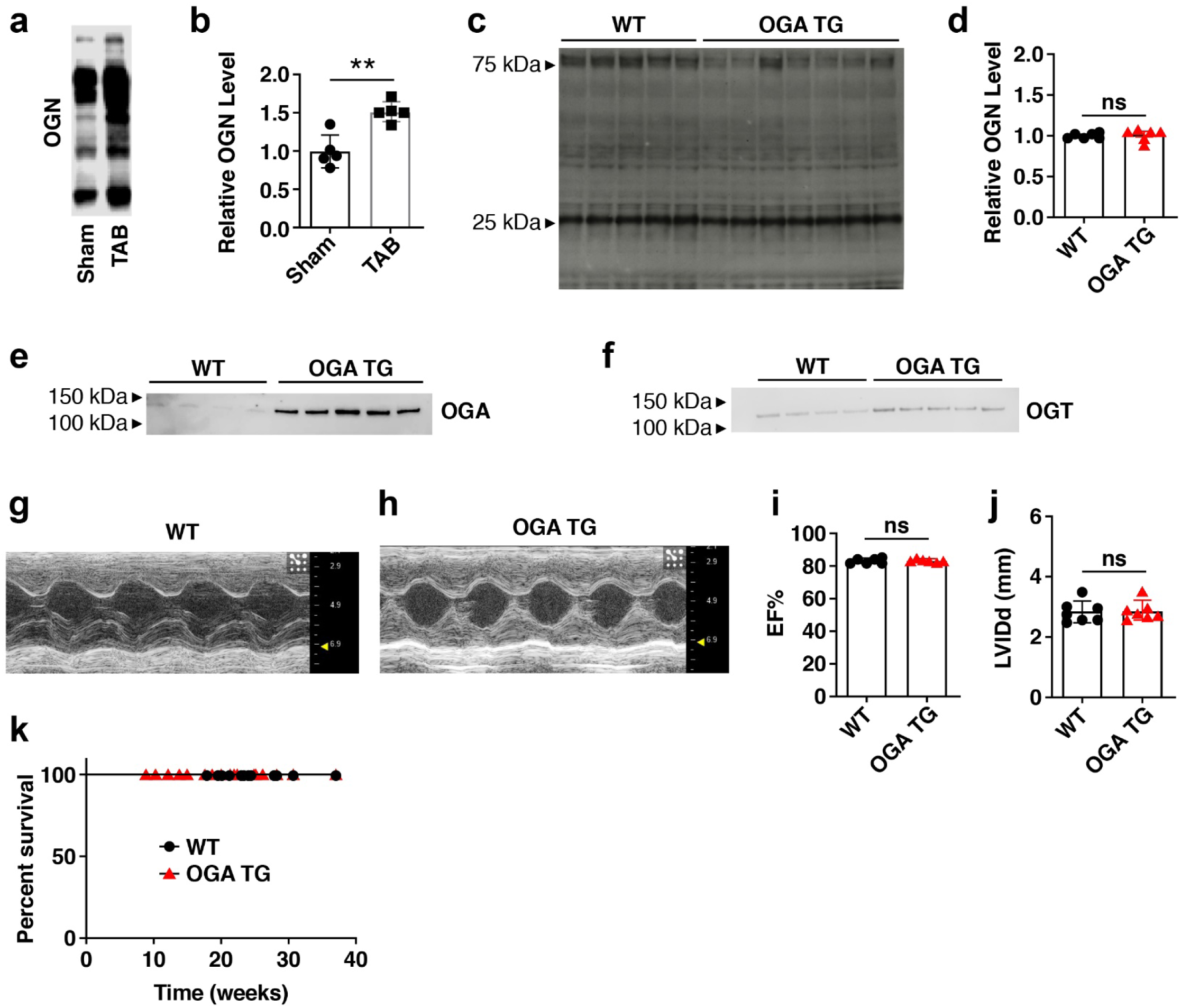
Myocardial OGA over-expression does not change *O*-GlcNAcylation nor cause cardiomyopathy. **a**. Representative western blot and **b.** summary data for total *O*-GlcNAc modified protein levels from whole heart lysates of 8-12 week old mice. Hearts were removed 14 days after transaortic banding (TAB) or sham surgery (n = 5 mice/group). **c.** Western blot of *O*-GlcNAcylation levels and **d.** summary data from cardiac lysates using WT (n=5) and OGA TG (n=7) mice. **e.** OGA and **f.** OGT protein expression from whole heart lysates from OGA TG (n=4) and WT (n=5) mice. **g.** and **h.** Example images of left ventricular M-mode echocardiograms from WT and OGA TG mice. Summary echocardiographic data for **i.** left ventricular ejection fraction (EF) and **j.** left ventricular end-diastolic internal diameter (LVIDd) acquired at 8-10 weeks of age (n= 7 WT, n=7 OGA TG). **k**. Kaplan-Meier survival analysis for OGA TG (n=11) and WT littermates (n=9). Data are represented as means ± SEM, significance was determined using the log rank (Mantel-Cox) test. (****P<0.0001, ***P<0.001, **P<0.01, *P<0.05, ns=not significant for all panels).

In order to test the effect of reducing myocardial *O-*GlcNAcylation under conditions of pathological stress, we developed OGA TG mice where OGA expression was under control of the α-myosin heavy chain promoter (Fig S1a, upper panel) (see Methods)^13^. The OGA TG mice were born in normal Mendelian ratios. The *O-*GlcNAcylation levels in OGA TG hearts were similar to WT hearts at baseline (Fig 1c and d). The myocardial OGA protein over-expression in OGA TG mice was increased several-fold over WT (Fig 1e), and OGA expression was restricted to the heart (Fig S1b). OGA activity was 20 fold increased in OGA TG compared to WT heart lysates (Fig S1c). OGA and OGT expression are coupled by mechanisms that include changes in transcription and splicing^14-16^ so we asked if the WT levels of *O-*GlcNAcylation in OGA TG hearts (Fig 1c and d), given elevated levels of OGA(Fig 1e) could be due to compensatory increased expression of OGT. We did find increased OGT expression (Fig 1f) and a trend toward increased OGT activity (Fig S1d) in OGA TG lysates, suggesting the basal myocardial *O-*GlcNAcylation level is sustained during OGA TG overexpression by increases in OGT expression. The OGA TG mice had modestly, but significantly increased heart weight/body weight ratios compared to WT littermate controls (Fig S2a), but no difference from WT mice in left ventricular function by echocardiography (Fig 1g-i), left ventricular dimensions (Fig 1j), or *Nppa* expression (Fig S2b), a marker of pathological hypertrophy^17^. The OGA TG mice did not show evidence of premature death during the first year of life (Fig 1k). We interpreted these findings to suggest myocardial OGA over-expression is well tolerated under basal conditions, perhaps because of a compensatory increase in OGT activity and expression that maintains *O-*GlcNAcylation homeostasis.

Elevated cardiac *O-*GlcNAcylation is associated with pathological hypertrophy and heart failure in animal models, and in patients with hypertension and aortic stenosis, conditions of increased left ventricular afterload^1^. Based on these associations, we next challenged OGA TG and WT littermate control mice with transaortic banding surgery(TAB) (Fig 2a). The OGA TG mice showed decreased myocardial *O*-GlcNAcylation (Fig 2b and c) after TAB. OGA TG mice were significantly protected against systolic dysfunction (Fig 2d), had thinner left ventricular posterior walls compared to WT mice (Fig 2e), and showed reduced expression of the myocardial hypertrophic reprogramming genes, *Nppa* and *Myh7* (Fig 2f and 2g), compared to WT littermate controls after transaortic banding surgery. Taken together, we interpreted these results to support a view that excessive *O-*GlcNAcylation can contribute to cardiomyopathy, and that increased OGA expression can reduce excessive *O*-GlcNAcylation and protect against cardiomyopathy.

**Fig 2.**
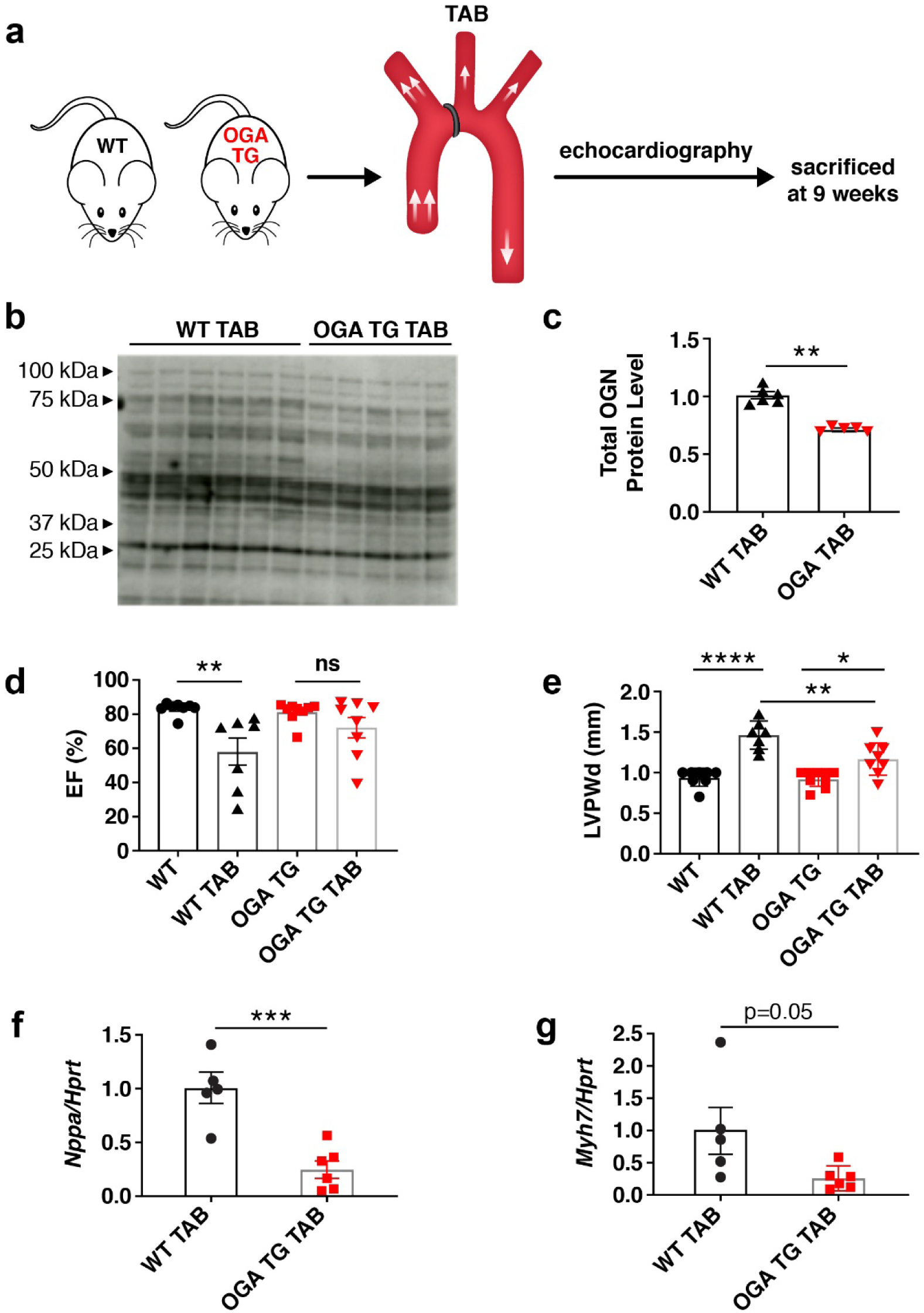
Myocardial-targeted OGA over-expression protects against left ventricular hypertrophy and contractile dysfunction after transaortic banding (TAB) surgery. **a**. Schematic of the TAB left ventricular pressure-overload model performed in 8-10 week old WT (n=6) and OGA TG (n=5) mice. **b**. Western blot and **c**. summary data of total *O*-GlcNAcylation levels from OGA TG (n=5) and WT (n=6) whole heart lysates 9 weeks after TAB. **d.** Left ventricular ejection fractions (EF) in OGA TG and WT littermate hearts 9 weeks after TAB surgery, and **e**. left ventricular posterior wall thickness measured at end-diastole (LVPWd) in WT (n=7) and OGA TG (n=8). **f**. Quantification of *Nppa* and **g.** *Myh7* mRNA expression normalized to Hypoxanthine Peroxidase Reductase Transferase (*Hprt*) in OGA TG (n=6) and WT (n=5) hearts 9 weeks after TAB. Data are represented as mean ± SEM. Significance was determined using a two-tailed Student’s *t* test or 1 way ANOVA with Tukey’s multiple comparison’s test, as appropriate (****P<0.0001, ***P<0.001, **P<0.01, *P<0.05, ns=not significant).

### Increased *O-*GlcNAcylation, dilated cardiomyopathy, arrhythmias, and premature death in OGT TG mice

Our findings up to this point showed reducing *O-*GlcNAcylation may be beneficial in cardiomyopathy due to transaortic banding surgery. We next used an orthogonal approach to assess the role of increased *O-*GlcNAcylation in cardiomyopathy by developing an OGT TG mouse model. OGT TG mice were designed for myocardial-targeted OGT over-expression, using the α-myosin heavy chain promoter, similar to our OGA TG mice (Fig S1a, lower panel) (see Methods). OGT TG and WT littermate mice were sacrificed at 8 weeks of age; OGT protein overexpression was exclusively localized to heart (Fig S3a). Cardiac *O-*GlcNAcylation (Fig 3a and b), and OGT protein expression were increased compared to WT littermates (Fig 3c). OGT activity in OGT TG was significantly increased compared to WT (Fig S1d). Cardiac OGA expression (Fig 3d) was very mildy increased over WT littermates, consistent with known reciprocal regulation of OGA and OGT expression^18^. The OGT TG mice developed dilated cardiomyopathy (Fig 3e and f) with reduced left ventricular ejection fraction (Fig 3g), and increased left ventricular diameter (Fig 3h). Left ventricular function began to decline after 6 weeks of age (Supplemental Fig S3b). By 8 weeks OGT TG mice had significantly increased left ventricular mass, assessed by echocardiography (Fig S3c), and by morphometric analysis (Fig S3d). These findings showed increased myocardial OGT was sufficient to cause cardiomyopathy.

**Fig 3.**
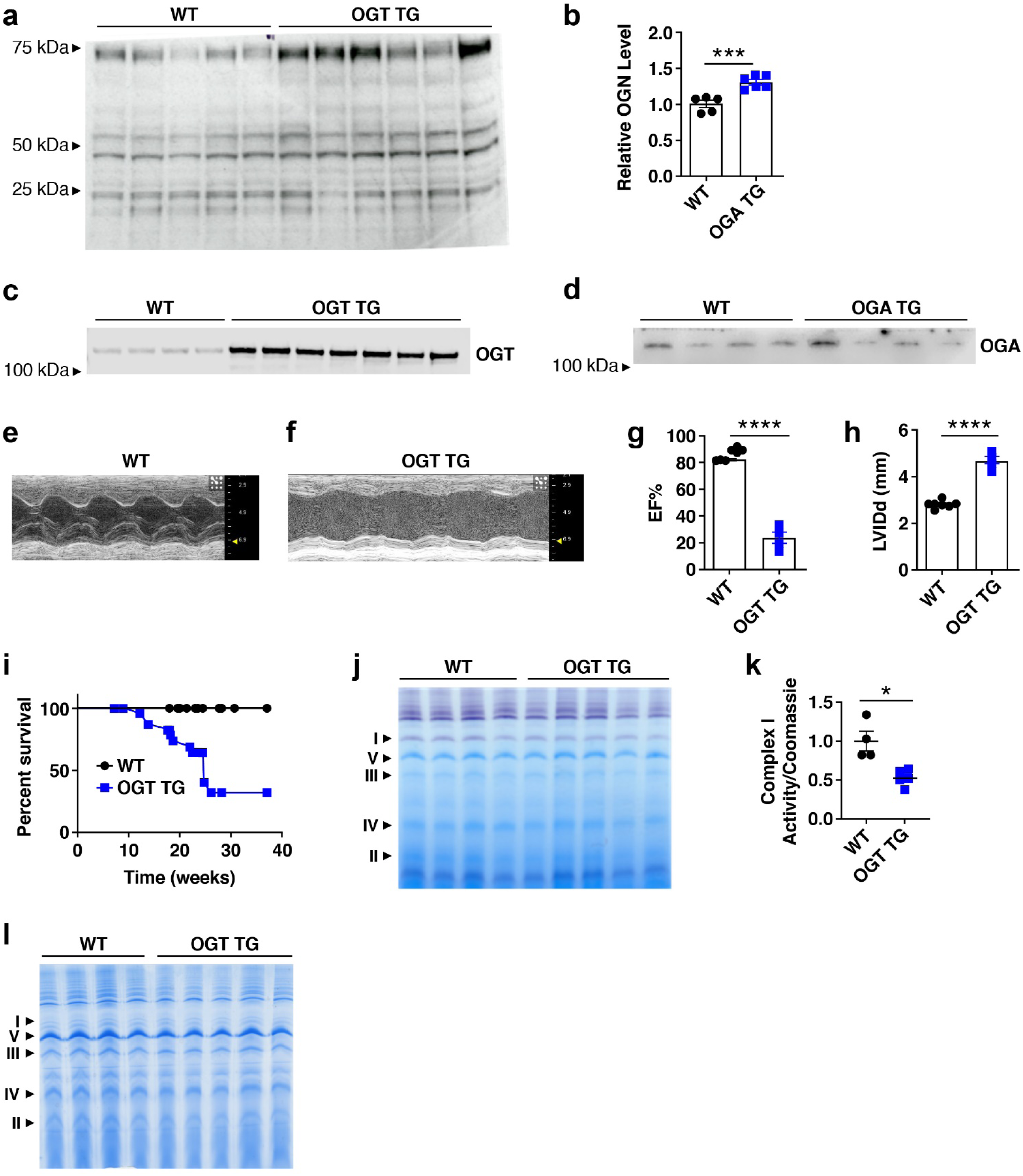
OGT TG mice have increased *O*-GlcNAcylation, dilated cardiomyopathy, and premature death. **a**. Western blot and **b.** summary data for total *O*-GlcNAcylation in cardiac lysates from OGT TG and WT littermates (n=5 WT, n=6 OGT TG hearts). **c.** OGT protein expression (n = 4 WT, n = 7 OGT TG) and **d**. OGA protein expression in cardiac lysates (n= 4 WT and n=4 OGT TG hearts). Example M-mode left ventricular echocardiograms from **e.** WT and **f.** OGT TG mice. **g**. Summary echocardiographic data for left ventricular ejection fraction (EF) and **h**. left ventricular internal diameter in diastole (LVIDd) in 8-12 week old mice. **i**. Kaplan-Meier survival analysis for OGT TG and WT littermate mice (n=9 WT, n=14 OGT TG). **j.** A blue native gel with WT (n=4) and OGT TG (n=5) mitochondrial isolates from heart stained for complex I activity. **k.** Summary data for complex I activity normalized to total mitochondrial protein expression. **l**. Total mitochondrial protein (1 heart/lane for panels j and l). Data are represented as mean ± SEM; significance was determined using a two-tailed Student’s *t* test (****P<0.0001, ***P<0.001, **P<0.01, *P<0.05, ns=not significant).

Sudden death is a major complication of many types of heart failure^19^ and OGT TG mice exhibited a striking pattern of premature mortality (Fig 3i). Arrhythmias are an important cause of premature mortality in heart failure^20, 21^, so we surgically implanted electrocardiographic monitors in OGT TG and WT littermate mice (20-22 weeks of age) to determine if OGT TG mice exhibited arrhythmias. We detected bradycardia, spontaneous ventricular tachycardia and ventricular fibrillation leading to death in OGT TG, but not in WT control mice (Fig S3e, left panel). The OGT TG mice exhibited a higher arrhythmia score (see Methods)^22^ (Fig S3e, right panel), confirming an increased arrhythmia burden in OGT TG compared to WT controls. These findings suggested that premature death in OGT TG mice was related, at least in part, to arrhythmias.

### Reduced oxidative phosphorylation in OGT TG cardiomyopathy

*O-*GlcNAcylation can modify thousands of proteins,^15, 16, 18, 23^ many of which could potentially contribute to cardiomyopathy. However, we focused on mitochondrial proteins because failing myocardium is marked by depressed energetics^24, 25^, and excessive *O-*GlcNAcylation can reduce oxidative phosphorylation^23^. We used blue native page gel electrophoresis ^26^ to analyze mitochondrial complex I, based on previous work highlighting complex I being a target for *O-*GlcNAcylation and altering mitochondrial respiration^11^. We evaluated the activity of complex I (Fig 3j and k), normalized to complex V protein expression (Fig 3l). We found marked reduction in complex I activity in OGT TG hearts (Fig 3j and k). These data show that OGT overexpression leads to increased *O-*GlcNAcylation, and loss of complex I expression and activity, suggesting OGT TG cardiomyopathy is due to depressed energetics.

### No evidence for increased myocardial death or deterioration of intracellular Ca^2+^ homeostasis in OGT TG hearts

Many types of acquired and genetic cardiomyopathies exhibit increased myocardial cell death^27^ and/or depressed intracellular Ca^2+^ concentration transients^28, 29^. Surprisingly, we did not detect differences in myocardial cell death (Fig S3f). These findings suggested that the loss of myocardial performance in OGT TG hearts was not due to loss of myocardium, or excessive myocardial scarring. Heart muscle cells contract and relax under control of intracellular Ca^2+^ concentration transients, which grade myofilament interactions to regulate muscle shortening^30^. We found that mechanically unloaded OGT TG ventricular myocytes had modest, but significantly reduced resting (diastolic) cytosolic Ca^2+^ concentrations (Fig S4a), and faster decay of peak (systolic) cytosolic Ca^2+^ (Fig S4b) compared to WT controls. OGT TG and WT ventricular myocytes had similar intracellular Ca^2+^ transient amplitudes (Fig S4c), and caffeine-releasable intracellular, sarcoplasmic reticulum, Ca^2+^ stores (Fig S4d). These findings suggested that OGT TG myocardium was not impaired due to defective intracellular Ca^2+^ homeostasis or paucity of sarcoplasmic reticulum Ca^2+^ reserve.

### Rescue of dilated cardiomyopathy and premature mortality in OGT TG and OGA TG interbred mice

We interpreted our findings in the OGT TG mice to suggest that excessive *O*-GlcNAcylation levels were a direct cause of cardiomyopathy. However, we also considered that OGT over-expression could have unanticipated pathological actions, independent of *O-*GlcNAcylation. To further test these possibilities, we interbred OGT TG and OGA TG mice. *O-*GlcNAcylation levels from OGT TG x OGA TG interbred hearts were similar to *O-*GlcNAcylation levels in OGA TG hearts, and significantly less than in OGT TG heart lysates (Fig 4a and b). Double transgenic mouse heart weights, adjusted for body weight, were less than OGT TG, and were slightly higher than WT littermates (Fig 4c). Echocardiography at 6-8 weeks of age revealed significant improvement in left ventricular ejection fraction (Fig 4d and e) and left ventricular dilation (Fig 4f). The rescue of *O-*GlcNAcylation levels, myocardial function and longevity in double transgenic mice occurred despite elevated levels of cardiac OGT (Fig 4g), and OGA protein expression (Fig 4h) compared to OGT TG hearts, suggesting that OGT TG cardiomyopathy was due to significantly elevated *O-*GlcNAcylation levels, rather than a non-specific consequence of transgenic protein over-expression. Remarkably, OGT TG x OGA TG mice were protected from the increased premature death seen in OGT TG mice (Fig 4i). Similar to our experiments in OGT TG mice (Fig 3j - l), we evaluated mitochondrial function using blue native gel activity assays for complex I. We found similar complex I activity in the interbred and WT mice (Fig 5a) normalized to complex V protein expression (Fig 5b). We then performed a complex I activity assay utilizing spectrophotometric quantification and confirmed that complex I activity was decreased, by approximately half, in OGT TG compared to WT and recovered to WT levels in double transgenic mice (Fig 5c). Finally we assessed oxygen consumption rates in isolated mitochondria in the four experimental groups (WT, OGT TG, OGA TG, OGT x OGA TG), utilizing the Seahorse XF96 Analyzer, and confirmed decreased complex I - linked respiration in OGT TG versus WT (Fig 5d). We observed a small, but statistically significant, difference between WT and OGA TG, and between OGA TG and OGT x OGA TG mitochondria in the absence of ADP (state 2, Fig 5e). However, the absolute difference between these groups was small, suggesting these differences may be of limited biological significance. In contrast, we measured larger differences in state 3 respiration (i.e. in the presence of ADP) between OGT TG, WT and OGT x OGA TG cardiac mitochondria (Fig 5f). We did not detect a difference in the state 3 oxygen consumption rates between WT and OGT TG x OGA TG mitochondria, suggesting that complex I activity is equivalent in these groups. Taken together, these results support a concept that OGT TG cardiomyopathy is driven, at least in part, as a consequence of impaired metabolism and energetics.

**Fig 4.**
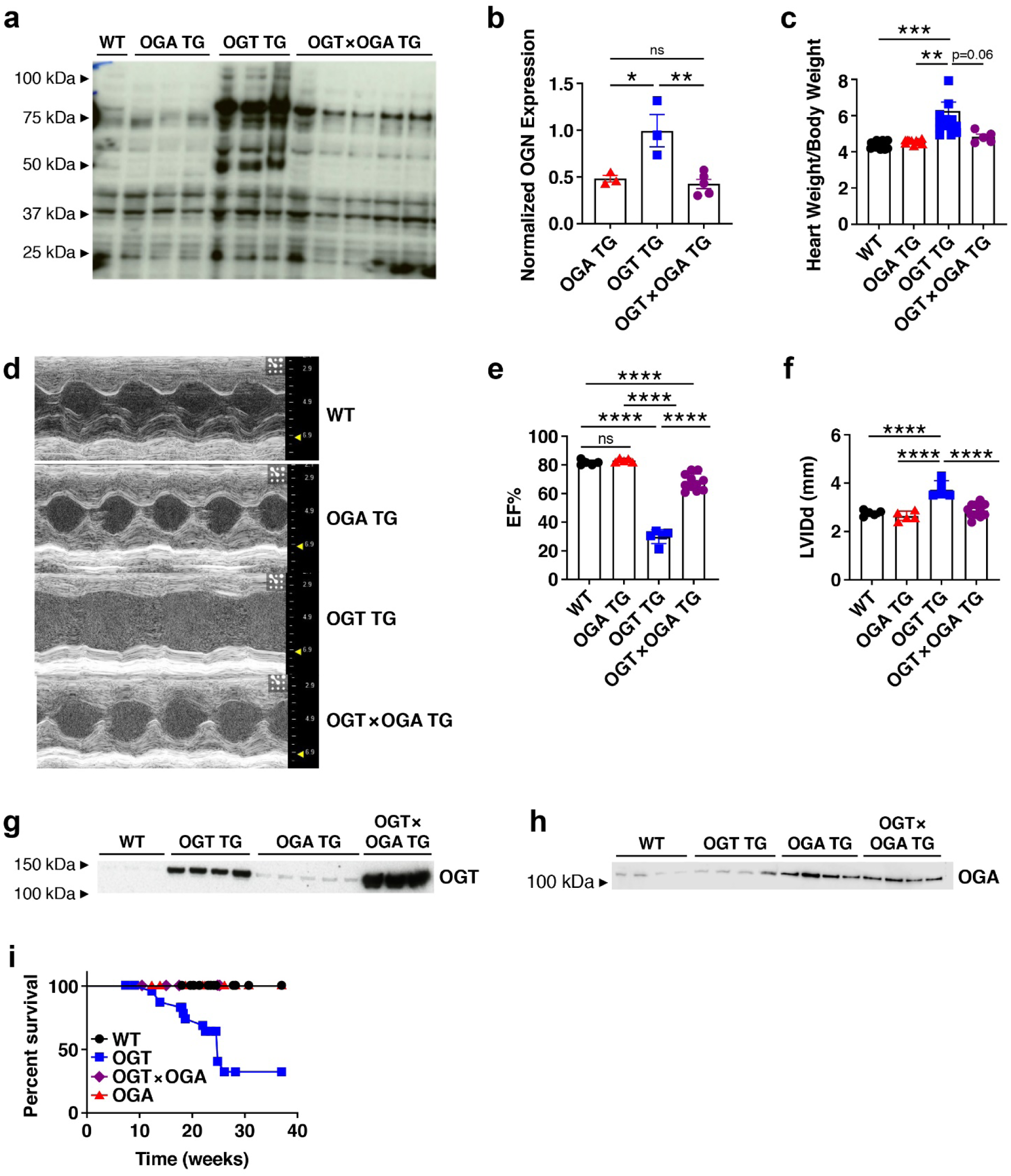
Rescue of OGT TG cardiomyopathy and premature mortality by OGA TG interbreeding. **a**. Western blot of total *O*-GlcNAcylation modified proteins in cardiac lysates from WT (n=1), OGA TG (n=3), OGT TG (n=3), and OGT x OGA TG (n=5) mice, and **b.** summary data. **c**. heart weight/body weight from WT (n=12), OGA TG (n =7), OGT TG (n = 15), OGT x OGA TG (n = 5) **d.** example images of left ventricular M-mode echocardiography of WT, OGA TG, OGT TG and OGT x OGA TG mice at 8-10 weeks of age. **e**. left ventricular ejection fraction (EF) and **f.** left ventricular internal diameter in diastole (LVIDd) **g.** western blot of OGT protein expression in hearts from WT (n=3), OGT TG (n=4), OGA TG (n=5), and OGT x OGA TG (n=3) mice **h**. OGA protein expression from WT (n=4), OGT TG (n=4), OGA TG (n=4), and OGT x OGA TG (n=4) hearts shown in **g. i**. Kaplan-Meier survival analysis for WT (n=9), OGA TG (n=11), OGT TG (n=14) and OGT TG x OGA TG mice (n=9). Data are represented as mean ± SEM, significance was determined using a two-tailed Student’s *t* test or log-rank test (survivorship). (****P<0.0001, ***P<0.001, **P<0.01, *P<0.05, ns=not significant for all panels).

**Fig 5.**
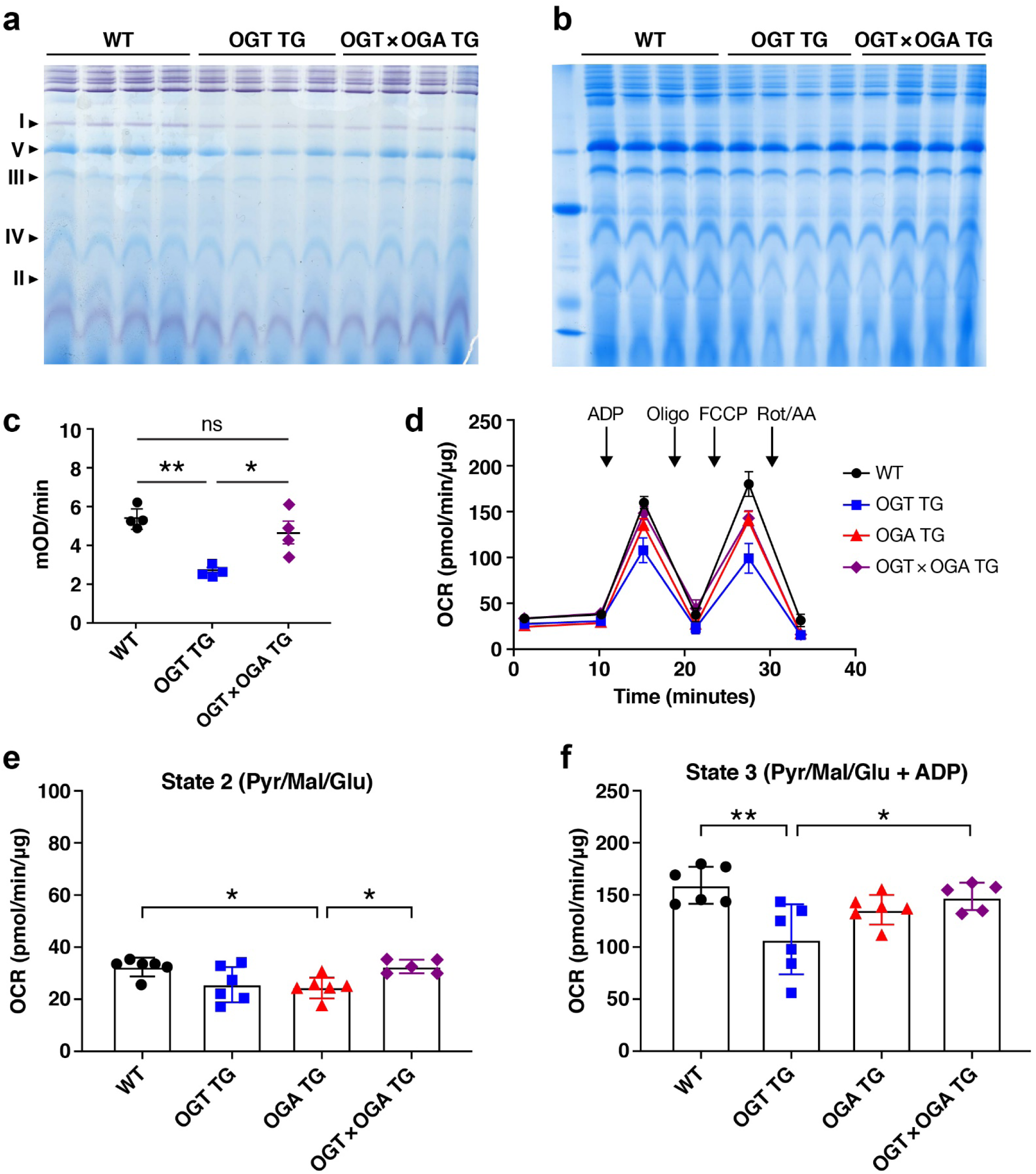
Reduced oxidative phosphorylation through impaired Complex I activity in OGT TG cardiomyopathy with rescued activity in OGT x OGA TG. Blue Native gel WT (n=4), OGT TG (n=4), OGT x OGA TG (n=4) stained for **a**. complex I activity, and **b.** Coomassie stain for total mitochondrial protein expression from heart (1 heart/lane), and **c**. Complex I activity spectrophotometer assay summary data. **d.** Measurement of isolated mitochondria OCR after sequential addition of ADP, oligomycin, FCCP and rotenone/antimycin A in the presence of substrates pyruvate, glutamate, and malate from all genotypes (WT, OGT TG, OGA TG n = 6, OGT x OGA TG n = 5). **e**. OCR measurement before ADP addition (the first time point in 5d) **f.** OCR measurement after ADP addition (the third time point in 5d). Data are represented as mean ± SEM, significance was determined using a 1 way ANOVA with Tukey’s multiple comparison’s test. (****P<0.0001, ***P<0.001, **P<0.01, *P<0.05, ns=not significant).

### Transcriptional reprogramming in OGT TG hearts

Given the known role of *O-*GlcNAcylation modification in modulating transcriptional pathways^15^ and the complexity of targets and pathways potentially under the influence of pathological *O-*GlcNAcylation, we hypothesized that multiple gene programs were affected in OGT TG cardiomyopathy. We compared polyA transcriptomes by RNA sequencing (RNA Seq) using hearts from age and gender matched WT, OGT TG, OGA TG and OGT TG x OGA TG interbred mice. Principle component analysis showed that each of these groups exhibited gene expression patterns that were more similar within than between groups (Fig 6a). In order to gain further insight into genes with the potential to drive *O-*GlcNAcylation cardiomyopathy, we focused on RNA Seq data gene sets that were significantly altered in OGT TG compared to WT hearts, and where these genes were repaired toward wild type expression in the OGT x OGA TG interbred hearts. We performed one way two*-*tailed Student’s *t*-test analyses comparing gene expression changes from CuffDiff data (multiple-test false discovery rate-adjusted q-value < 0.05) between the WT versus OGT TG, and the OGT TG versus OGT x OGA TG groups. We found 2813 genes were significantly changed in the OGT TG versus WT hearts, and 1802 genes were significantly altered in the OGT x OGA versus OGT TG hearts, while 1798 individual genes showed significant expression differences between the WT and the OGT x OGA TG hearts. We applied QIAGEN Ingenuity Pathway Analysis and identified the top 5 significant canonical pathways and biological functions in OGT TG and OGT x OGA TG hearts using a regulation z-score and an overlap p-value (Fisher’s exact test; p < 0.05). The most prominent functional pathways identified included inhibition of oxidative phosphorylation pathways (Fig 6b) in the OGT TG hearts compared to hearts from OGT x OGA interbred mice, and activation of pathways involved in inflammation, inflammatory cell signalling and transcription (Fig S5a).

**Fig 6.**
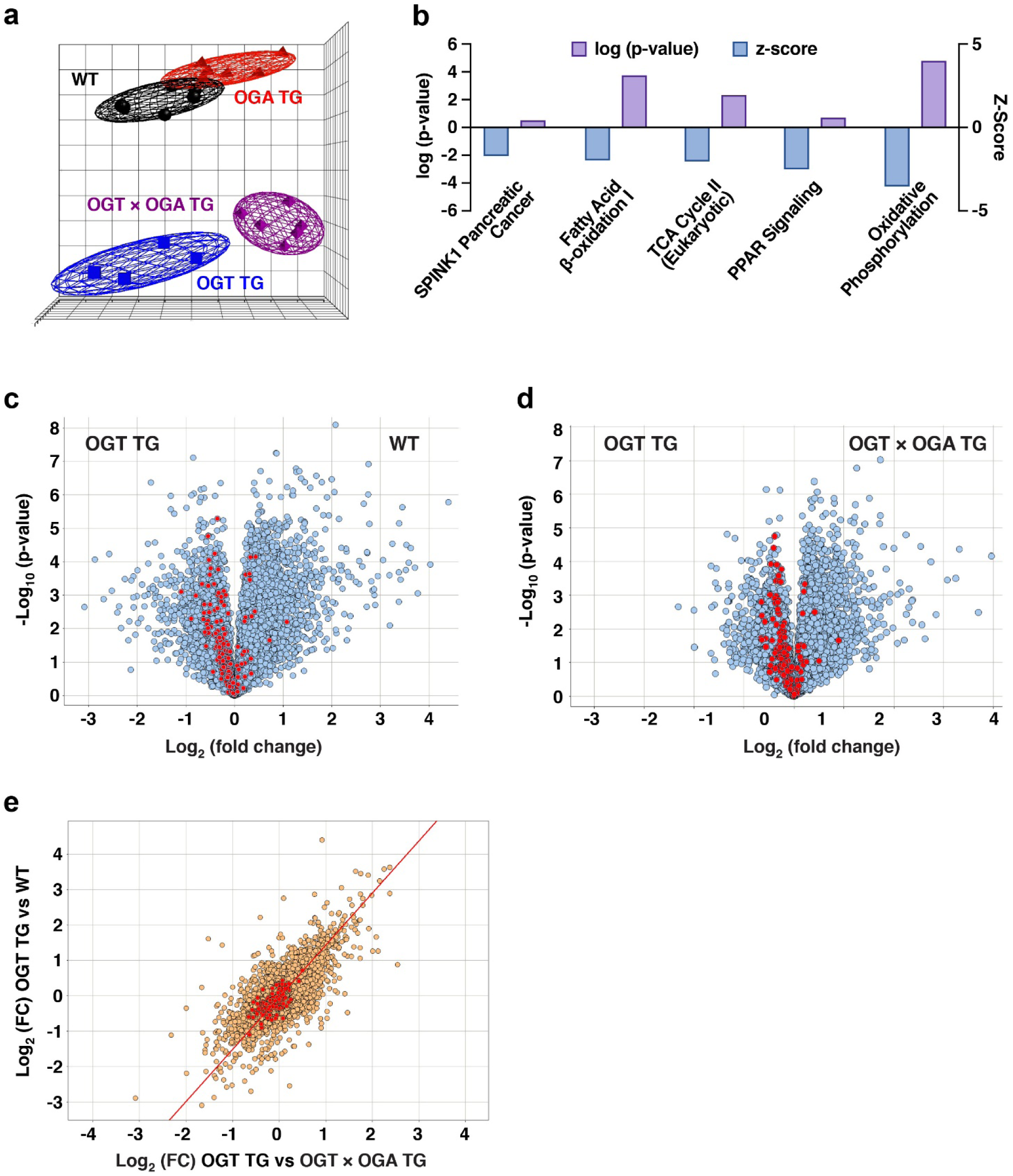
Metabolic gene expression defects in OGT TG mice are recovered by OGA TG interbreeding. **a.** Principal component analysis of RNA Seq data from WT, OGT TG, OGA TG and OGT x OGA TG hearts (n=6 in all groups, M=F) demonstrating clustering by similarity of transcriptome. **b.** The top 5 significant canonical pathways and biological functions identified by Ingenuity Pathway analysis with a regulation z-score and an overlap p-value (Fisher’s exact test; p < 0.05) for the comparison of significant differentially expressed genes in the dataset. The Z-score represents the observed up or down regulation compared to known changes that are either activating or inhibiting, as derived from the literature and compiled in the Ingenuity^®^ Knowledge Base. Pathways represented here are overall inhibited. Volcano plots representing gene set enrichment analysis of hallmark genes (as derived from the Molecular Signature Database) for oxidative phosphorylation comparing **c.** OGT TG versus WT and **d.** expression of oxidative phosphorylation genes in OGT x OGA TG versus OGT TG All genes are represented in grey and hallmark oxidative phosphorylation genes are represented in red. **e**. Q-q plot comparing overlapping genes between OGT TG versus WT and OGT TG versus OGT x OGA TG (all genes in gold and hallmark gene set oxidative phosphorylation genes in red, FC = fold change).

The prominent changes exhibited in oxidative phosphorylation pathways aligns with our data from blue native gels and complex I activity assays (Fig 5) that suggested impairment in mitochondrial energetics, with subsequent recovery in the interbred OGT TG x OGA TG mice. We compared the hallmark oxidative phosphorylation gene set defined by Liberzon et al^11^. to genes identified by our RNA sequencing study, and identified 177 genes out of 200 genes listed in this gene set. We found that the majority of these genes (highlighted in red) are downregulated in OGT TG hearts compared to both WT hearts (Fig. 6c) and OGT x OGA TG (Fig. 6d) hearts, suggesting that OGT TG hearts are deficient in expression of oxidative phosphorylation genes, and OGT x OGA TG interbreeding recovered their expression towards WT levels. This observation was further supported by hierarchical clustering analysis (Fig S5b) and Q-Q plot analysis (Fig 6e.). These data appeared to confirm our experimental findings, and support published work^31^, highlighting defective mitochondrial energetics in response to excessive *O-*GlcNAcylation.

## Discussion

The association between increased *O-*GlcNAcylation with diverse forms of cardiac stress is well known^32-34^. However, to our knowledge, no direct causal relationship has been established between increased *O-*GlcNAcylation and cardiomyopathy. Notably, mouse models with near elimination of myocardial OGT exhibited exaggerated responses to injury^2^. We interpret this important finding to indicate some increase in *O*-GlcNAcylation, likely within an acute or subacute timeframe, is required for optimal cardiac responses to stress. However, our results show therapeutic benefit from modest reduction of *O-*GlcNAcylation in OGA TG mice after aortic banding, and severe cardiomyopathy, heart failure and sudden death resulting from massive and chronic *O*-GlcNAcylation increases in OGT TG mice. Collectively, we interpret these data to strongly suggest excessive myocardial *O-*GlcNAcylation contributes to cardiomyopathy. In contrast to our new data, most work supporting a connection between excessive *O*-GlcNAcylation and cardiomyopathy has focused on hyperglycemic conditions, including in models of type I diabetes^23^. Our study provides new evidence that exposure to chronic, excessive *O-*GlcNAcylation is sufficient to cause dilated cardiomyopathy and premature death, even in the absence of hyperglycemia, diabetes or metabolic disease. This finding may be broadly important because excessive *O-*GlcNAcylation could add to other established cardiomyopathy mechanisms, and because it suggests that reversing or preventing excessive myocardial *O-*GlcNAcylation could be an innovative and effective therapeutic strategy for cardiomyopathy and heart failure.

We recognize that transgenic models may imperfectly represent the pathological potential of pathways linked to protein over-expression, and this caveat could apply to the OGT TG mouse. However, the findings that OGT TG cardiomyopathy was rescued by interbreeding with OGA TG mice, resulting in very high levels of transgenic protein over-expression, strongly suggests that OGT TG cardiomyopathy was a specific consequence of excessive OGT activity. It is possible that *O-*GlcNAcylation modified proteins are different in OGT TG hearts compared to WT hearts with increased *O*-GlcNAcylation due to pathological stress. The OGA TG mice have very mild myocardial hypertrophy, but no other measured phenotypic differences compared to WT littermate controls. Our finding that OGA TG mouse hearts had less stress induced *O*-GlcNAcylation and were resistant to transaortic banding surgery suggests that transgenic overexpression of OGA is well tolerated, and is capable of reversing pathological *O-*GlcNAcylation modifications.

Our RNA Seq studies showed a wide variety of transcripts were affected by increased *O-*GlcNAcylation through OGT overexpression. Expression of genes involved in metabolism was reordered toward WT levels in OGT TG x OGA TG hearts. The role of depressed energetics and deranged metabolism is a consistent finding in dilated cardiomyopathy in patients^10, 25, 27^ furthermore, excessive *O-*GlcNAcylation is associated with defects in oxidative phosphorylation, in part, by actions in mitochondria^11, 23, 31, 35, 36^. We found reduced expression of a number of genes encoding complex I proteins, loss of complex I proteins, and decreased complex I activity in OGT TG hearts. Complex I gene expression, complex I proteins and complex I activity were remodeled toward WT levels in the OGT TG x OGA TG interbred hearts, suggesting that defective metabolism contributes to OGT TG cardiomyopathy.

In contrast to myocardial OGT overexpression, we found that massive myocardial OGA overexpression did not cause cardiomyopathy, and that OGA TG hearts were partly resistant to transaortic banding induced cardiomyopathy. These results suggested to us that reducing chronic *O-*GlcNAcylation elevation by OGA activation could be a new therapeutic strategy for cardiomyopathy. The cardiomyopathy phenotypes in OGT TG hearts were notable for the absence of increased fibrosis, or myocardial death, features often associated with irreversible disease^37-39^. Thus, it may be that *O-*GlcNAcylation contributes to heart disease by mechanisms, including depressed energetics, that are reversible. Given the public health consequences of cardiomyopathy and heart failure despite current treatments, the quest for improved therapeutic options remains an important goal for patients. Our findings suggest that targeting excessive *O*-GlcNAcylation could reduce cardiomyopathy and heart failure

## Acknowledgements

Grateful thanks to Teresa Ruggle at the University of Iowa for help with figure formatting.

