## Supplemental Material for "Excessive *O* - GlcNAcylation causes heart failure and sudden death"

**Supplemental Expanded Methods**

*Transaortic banding surgery*

The mouse was intubated and mechanically ventilated at 120 breaths per minute with a tidal volume of 200μl. The aortic arch was approached via minimal parasternal incision. Once the arch was visible, a small hemoclip (Weck Horizon) was placed around the transverse aorta, to the diameter of a 27g needle, between the innominate artery and the left carotid artery. After the clip was placed, the isoflurane was gradually decreased as the chest was closed. The musculature was closed with 5-0 vicryl and skin closed with 5-0 silk. The isoflurane was turned off and the mouse allowed to regain respiration. Within a few minutes, the mice were extubated and allowed to regain consciousness.

*Telemetry*

Telemeters were implanted in wild type and OGT TG mice at 18-20 weeks of age. General inhalation anesthesia was induced by placing the animal in the induction chamber and introducing the volatile anesthetic agent (sevoflurane 8% or isoflurane 5% in pure oxygen 600 ml/min). Anesthesia was maintained by spontaneous breathing (sevoflurane 3-4% or isoflurane 1.5-3% in pure oxygen at a flow rate of 600 mL/min).The skin of the anterior neck and abdominal region was disinfected with 70% ethanol and a 1- to 1.5-cm-long incision in the skin was made from the lower thorax along the midline to the abdomen. The negative (white/colorless) lead was tunneled subcutaneously from the thorax to the neck, where a small incision (≤0.5 cm) was made in the longitudinal direction. The wire loop is fixed between the muscles located to the right of the trachea, using vicryl sutures. The wound in the neck was closed with absorbable sutures (VICRYL 6-0, Ethicon, Norderstedt, Germany) in layers. The abdominal wall was opened at the linea alba and the body of the telemetric transmitter was placed into the abdominal cavity. The wire loop of the positive (red) electrode was sutured to the xiphoid process with silk. The skin of the abdominal region was restored with staples (Precise, 3 M Health Care, St. Paul, MN, USA).

After completion of surgery and anesthesia, 0.1 mg/kg of buprenorphine (Temgesic, Essex Chemie AG, Lucerne, Switzerland) was administered subcutaneously for pain treatment. During the recovery period of 4-10 days monitoring of general condition and body weight, as well as food and water consumption, was performed once daily according to the general condition and health monitoring data sheet for 10 days post-operatively.

*Arrhythmia Screening*

Arrhythmias were categorized into 5 groups[1] and assigned the following point values: no arrhythmias, 0 points; premature atrial or ventricular beats, 1 point; supraventricular tachycardia or paired premature ventricular beats, 2 points; bigeminal or trigeminal premature ventricular beats or nonsustained ventricular tachycardia (≥3 consecutive premature ventricular beats), 3 points; and sustained ventricular tachycardia (>10 consecutive premature ventricular beats) or polymorphic ventricular tachycardia, 4 points.

*Transthoracic echocardiography*

We recorded transthoracic echocardiograms in conscious mice. Images were acquired and analyzed by an operator blinded to mouse genotype and treatment. Unanesthetized mice were used for echocardiography. The anterior chest was shaved and warmed gel was applied. The mouse was grasped gently by the nape of the neck and positioned in the left lateral position. A 30-MHz linear array transducer was applied to the chest to obtain cardiac images. The transducer was coupled to a Vevo 2100 imager (FUJIFILM VisualSonics). Images of the short and long axis were obtained with a frame rate of ~180–250 Hz. All image analysis was performed offline using Vevo 2100 analysis software (Version 1.5). Endocardial and epicardial borders were traced on the short axis view in diastole and systole. The left ventricular length was measured from endocardial and epicardial borders to the left ventricular outflow tract in systole and diastole. The biplane area-length method was then performed to calculate left ventricular mass and ejection fraction.The left ventricle end-diastolic and end-systolic ventricular volumes (EDV, ESV), and the percent ejection fraction (EF) were estimated using the Simpson's method from the apical two chamber view of the heart[2]. Measurement of EDV, ESV were manually traced according the American Society of Echocardiography. The EF was automatically calculated by the system building software using the following formula:

$$EF(\%) = \frac{EDV - ESV}{EDV} * 100$$

The M-mode echocardiogram was acquired from the parasternal short axes view of the left ventricle (LV) at the mid-papillary muscles level and at sweep speed of 200 mm/sec. From this view left ventricle mass (LV mass) is calculated from inter-ventricular septal thickness at end of diastole (IVSD), LV chamber diameter at end of diastole (LVEDD), posterior wall thickness at end of diastole (PWTED) measurements using the following formula where 1.055 is the specific gravity of the myocardium.

$$LV\ Mass\ (mg) = 1.055 * \left((LVSD + LVEDD + PWTED) * 3 - (LVEDD) * 3\right)$$

For all the cardiac function studies, the observer was blinded to the experimental groups.

***Protein Studies***

*Western blot*

Heart lysates were prepared in buffer containing 0.1% Triton-X, PUGNAc, Thiamet G, protease and phosphatase inhibitors. Heart lysate or cytoplasm and mitochondria fractions were loaded into 4-12% gels (Bio-Rad). Proteins were transferred to nitrocellulose membrane using Turbo-blotter (Bio-Rad). Primary antibodies *O*-GlcNAcylation (CTD 110.6, kind gift from Gerald Hart), OGT (Abcam), OGA (Abcam), Myc (Rockland), GAPDH (Cell Signaling), and α-actinin (Sigma), were incubated with the membrane overnight at 4□C. Blots probed using CTD 110.6 were blocked with 5% BSA for one hour and incubated overnight at 4□C, murine IgM-HRP was used as a secondary for CTD 110.6 blots and incubated at room temperature for one hour. Blots were imaged using an Odyssey Fc Imager (Licor) except CTD 110.6 blots were imaged using film (Fujifilm). Densitometry was performed on non-saturated exposures and analyzed using the ImageJ software (NIH). The relative signal was captured from the entire lane when analyzing probes that yielded multiple bands. Signals were normalized to Coomassie stain for total protein to control for loading differences.

*OGA Activity Assay*

Lysates in lysis buffer (50mM HEPES, 0.1% NP-50) at an equal concentration (2 mg/ml) were desalted into desalting buffer (20 mM Tris, pH 7.8, 20% (v/v) glycerol containing protease and phosphatase inhibitors) using Zeba spin desalting columns (7,000 molecular weight cutoff; Thermo Fisher Scientific) and then the protein concentration was re-assessed.

The activity of OGA was assessed in total cell lysate (30 µg) of 4 biological replicates (6 technical replicates) using 1 mM 4MU-GlcNAc fluorescent substrate in OGA reaction mixture (Final, 100 mM sodium cacodylate, pH 6.4 (50 mM in Fig. 2C), 2mg/ml BSA, 100mM GalNAc) in a black flat/clear-bottomed 96-well plate. β-N-Acetylhexosaminidase F (New England Biolabs, Ipswich, MA) diluted in desalting buffer was used as a positive control (5 units/well). Assays were quenched with glycine, pH 10.75 (final concentration, 150 mM in 50-µl assays), and the fluorescence intensity was measured using an Spectramax I3X Multimode Microplate detection platform (Molecular Devices, San Jose, CA; excitation 360 nm and emission 460 nm). Activity was converted to picomoles/min/mg using a standard curve of free 4MU.

*OGT Activity Assay*

Lysates in lysis buffer (50mM HEPES, 0.1% NP-50) at an equal concentration (2mg/mL) were desalted into desalting buffer (20 mM Tris, pH 7.8, 20% (v/v) glycerol) using Zeba spin desalting columns (7,000 molecular weight cutoff; Thermo Fisher Scientific) and the protein concentration was re-assessed. In a clear round-bottomed 96-well plate, 30 µg of cell lysate (4 biological replicates) was incubated (1 h, 25 °C) in triplicate with 0.5 µCi of [3H]UDP-GlcNAc (ART 0128; American Radiolabeled Chemicals, St. Louis, MO; specific activity, 60 Ci/mmol), 1 mM casein kinase II (CKII) acceptor peptide (PGGSTPVSSANMM; The Johns Hopkins University School of Medicine Synthesis and Sequencing Facility), 2.5 units of calf intestinal alkaline phosphatase (New England

Biolabs), 0.25 mM 5′-AMP (Sigma), and OGT assay buffer (100 mM sodium cacodylate, pH 6.4, 2mg/ml BSA). Recombinant His6-OGT was purified in-house on nickel-nitrilotriacetic acid-agarose (Qiagen, Venlo, The Netherlands), and used as a positive control (0.5 µg/well). Assays were quenched with 37.5 mM formate, 750 mM NaCl (final concentration). Samples were loaded onto a Strata $C_{18}$ 96-well plate (25 mg/well; 8E-S001-CGB; Phenomenex) activated with 100% methanol and equilibrated in 50 mM formate, 1 M NaCl (three times with 2 ml each). The $C_{18}$ plate was subsequently washed with 50 mM formate with 1 M NaCl, water, and 50 mM formate (two times with 2 ml each). The reaction product was eluted from the column with 100% methanol (2 ml), and the incorporation of radiolabeled GlcNAc was assessed by liquid scintillation counting (Beckman Coulter Inc., Indianapolis, IN). Activity was normalized by subtracting the counts arising from the average of triplicate samples incubated without CKII acceptor peptide. Disintergrations per minute were used to calculate the pmoles of GlcNAc added per minute per mg of lysate.

***Metabolism studies***

*Mitochondrial isolation*

Mitochondria were isolated using homogenization method on ice as previously described[3]. In brief, the heart was minced and homogenized in ice-cold isolation buffer (75 mM sucrose, 225 mM mannitol, 1 mM EGTA, and 0.2% fatty acid free BSA, pH 7.4) using a Potter-Elvehjem glass homogenizer. The homogenates were centrifuged for 10 min at 500Xg. The resulting supernatants were centrifuged for 10 min at 10,000Xg. The mitochondrial pellets were washed twice in isolation buffer and centrifuged at 7,700Xg for 6 min each. Mitochondrial pellets were suspended in a small amount of isolation buffer to ~5mg/ml. The protein concentration was determined by using the BCA Assay kit (Thermo Fisher Scientific).

*Blue Native Gel Electrophoresis*

Mitochondrial isolates from heart tissue lysates were prepared according to the NativePAGE Sample prep kit (ThermoFisher) (8g/g: digitonin/protein). The mitochondrial resuspensions 3-12% Bis-Tris Native Page gels (ThermoFisher). Gels were run using the NativePAGE Running Buffer kit (ThermoFisher). Gels were either stained using the Coomassie R-250 Staining kit (ThermoFisher) or used for in-gel activity. For this, the gel was equilibrated in 5 mM Tris HCl, pH 7.4 for 10 min and then developed in complex I activity solution (5 mM Tris pH7.4, 2.5mg/ml NBT, 0.1mg/ml NADH). The reaction was terminated by soaking the gel in 10% acetic acid. Gel were imaged using Epson Perfection V800 Photo scanner.

*Complex I Enzyme Activity Microplate Assay*

The activity of mitochondrial complex I was measured using the complex I enzymatic activity microplate assay kit (Abcam-MitoSciences, Cambridge, MA, USA) according to the manufacturer’s instructions. Enzymatic activity was monitored at 450 nm on a microplate reader (ThermoFisher Scientific, Waltham, MA, USA) by following the oxidation of NADH to oxidized nicotinamide adenine dinucleotide (NAD+). Values were normalized to the total protein concentration of the same sample.

*Seahorse Mitochondrial Bioenergetics Assay*

The isolated mitochondria were suspended in a potassium based buffer (2µg in 25µl of 137mM KCl, 2mM KH2PO4, 0.5mM EGTA, 2.5mM MgCl2, 20mM HEPES, pH 7.2) and loaded onto the XFe/XF96 cell culture microplate (Agilent) and centrifuged at 2000g for

20 minutes at 4°C. The substrate (10mM pyruvate, 2mM glutamate, 2mM malate with 0.2% (w/v) BSA in potassium based buffer) was added into the cell culture microplate before the experiment. After the sensor calibration and baseline OCR measurements, the port injections were as follows: port A, ADP (2mM, final); port B, oligomycin (2 μM, final); port C, FCCP (2 μM, final); and port D antimycin A and rotenone (1μM, final). In each port injection, the mixing time is 1 minute, measuring time 3 minutes and followed by another 1 minute of mixing.

***Cell and tissue studies***

*Adult ventricular myocyte isolation*

Adult ventricular myocytes were isolated as previously described[4]. Briefly, mice (8-12 weeks, either gender) were anesthetized by Avertin injection. Hearts were rapidly excised and placed in ice cold nominally $Ca^{2+}$ free HEPES-buffered Tyrode's solution. The aorta was cannulated, and the heart was perfused in a retrograde fashion with a nominally $Ca^{2+}$ free perfusate for 5 min at 37°C. This was followed by a 15-min perfusion with collagenase-containing nominally $Ca^{2+}$ free solution. Final perfusion was with collagenase-containing low $Ca^{2+}$ (0.2 mM) solution. The LV and septum were cut away, coarsely minced and placed in a beaker containing low $Ca^{2+}$ solution with 1% (w/v) BSA at 37°C. Myocytes were dispersed by gentle agitation, collected in serial aliquots and then maintained in standard saline solution containing 1.8 mM $Ca^{2+}$.

*Cytosolic $Ca^{2+}$ Measurements*

Cytosolic $Ca^{2+}$ levels were recorded from Fura-2–loaded ventricular myocytes as previously described[4]. Briefly, single isolated ventricular myocytes were loaded with 4 μM Fura-2 acetoxymethyl (AM) for 30 min and then perfused with Tyrode's solution for

30 min to de-esterify the Fura-2 AM. After placement on a recording chamber, the cells were perfused in bath solution comprised (mM): 137 NaCl, 10 Hepes, 10 glucose, 1.8 $CaCl_2$, 0.5 $MgCl_2$, and 25 CsCl; pH was adjusted to 7.4 with NaOH, at 35 ± 0.5 °C. The pipette (intracellular) solution comprised (mM): 120 CsCl, 10 Hepes, 20 TEA chloride, 1.0 $MgCl_2$, 0.05 $CaCl_2$, 10 glucose; the pH was adjusted to 7.2 with 1.0 N CsOH. Adenosine 5'-triphosphate disodium salt hydrate 5 mM was added to pipette solution when needed. Myocytes were stimulated at 1, 3 and 5Hz using voltage protocol holding at −80 mV and step to 0 mV for 100 ms from a prepulse of 50 ms at −50 mV. The cytosolic $Ca^{2+}$ transients were measured from cells excited at wavelengths of 340 and 380 nm and imaged with a 510-nm long-pass filter.

*TUNEL staining*

Cryosections (10μM) of ventricular tissue were fixed in 4% paraformaldehyde and stained with In-situ Cell Death Detection kit (Roche). Sections were imaged on a laser-scanning confocal microscope (LSM 510, Carl Zeiss). TUNEL positive and total nuclei were counted from 5 images per sample, and the averages reported.

***Gene Expression and Analysis***

*RNA-Seq and data analysis*

Whole hearts were isolated from WT, OGT TG, OGA TG and OGT x OGA TG (n = 3 male and n=3 female in each group) and RNA was prepared for subsequent RNA-Seq analysis to document transcriptome changes associated with the transgenic genotype and potentially identify differentially expressed genes.

Transcriptomic data collected by RNA-seq were analyzed to evaluate the genes that are transcribed in each sample and condition, their expression levels, and the differences in expression levels between experiment conditions. Following quality review using the software Fastqc**,** the reads were mapped to the mouse genome (version mm10) augmented with the human OGT and OGA cDNA sequences, using the alignment tool Tophat2 v.2.1.0[5]**.** The aligned reads were assembled with StringTie v.1.3.4c[6] to create partial gene and transcript models (transfrags). Transfrags from all samples were further merged with StringTie (ST)-merge and mapped to the mouse RefSeq gene models, to create FPKM files with a unified set of gene annotations for differential analyses. Gene and transcript expression levels were computed, and differentially expressed genes and transcripts were determined with the tools Cuffdiff2 v.2.2.1[7] **.** Lastly, differentially spliced events (exon skipping, mutually exclusive exons, alternative 5' and 3' splice sites and retained introns) were determined with the program rMATS v.3.2.5[8].

The FPKM files were converted to log notation and quantile normalized to minimize technical variation, using the Partek Genomics Suite v7.0 platform (Partek Inc, St. Louis MO). PCA plots demonstrated good within-class clustering, so the three biological classes were compared using the one way two-tailed t-test to determine each gene's relative expression level, as fold change, and those fold changes' statistical significance as q-values.

We used the downstream functional annotation platform, Ingenuity Pathway Analysis (QIAGEN Inc., https://www.qiagenbioinformatics.com/products/ingenuitypathway-analysis) to compare differentially-expressed genes with those that were not differentially expressed for each comparison, where a log2 fold change greater than

2SD defined differential expression. The enrichment score of each functional annotation cluster is the –log10 of the Fisher's exact test p-value of each annotation or pathway term (e.g. an enrichment score of >1.3 is equivalent to non-log scale $p < 0.05$). Therefore the higher the enrichment score, the more likely these annotations have a biologically relevant role. Annotation terms were manually selected from the list of representative terms generated within each cluster.

*Statistical Analysis*

Statistical analyses were performed using Graph Pad Prism 7 software. Sample size and information about statistical tests are reported in the figure legends. Data are presented as mean ± SEM.  Pairwise comparisons were performed using a two-tailed Student's *t* test.  For experiments with more than 2 groups, data were analyzed by 1 way ANOVA followed by Tukey's post-hoc multiple comparisons test. For Kaplan-Meier survival analysis data are represented means ± SEM, significance was determined using the log rank (Mantel-Cox) test.

### Supplemental Figures

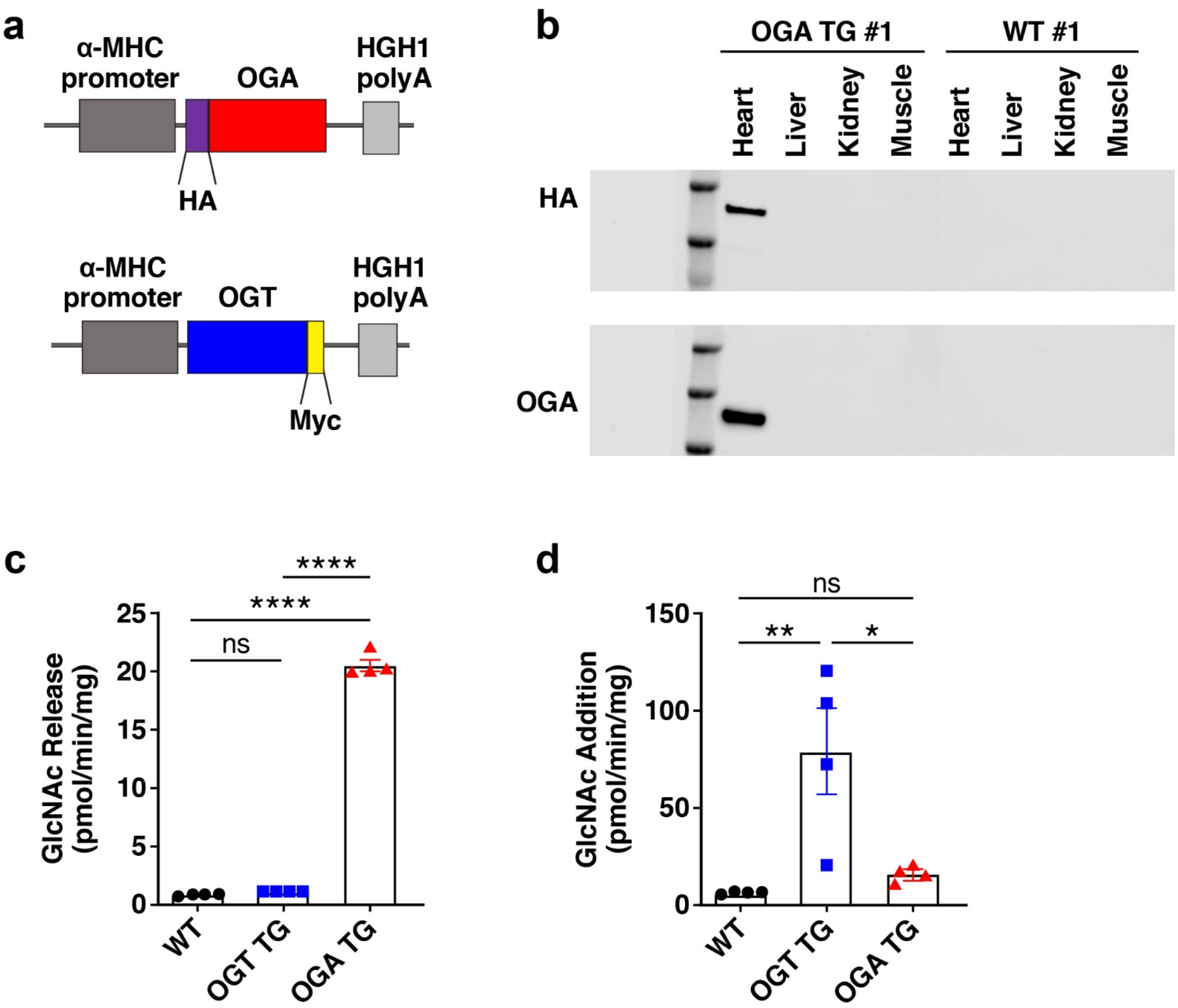


**Fig S1 - Supplemental Figure 1**

**Overexpression of OGA is cardiac specific**

**a.** Schematic of α-MHC promoter driven OGA transgenic construct with an HA epitope marker (upper panel) and OGT transgenic construct with Myc tag (lower panel)**. b.** Representative Western blot for HA-OGA transgene expression in heart, liver, kidney and muscle. Protein isolated from WT and OGA TG tissues (WT n=3, OGA TG n =3, OGT TG=3). **c**. OGA activity assay measuring GlcNAc release in WT (n = 4), OGT TG (n = 4) and OGA TG (n = 4) mouse hearts at 8-10 weeks. **d.** OGT activity assay measuring *O*-GlcNAc addition in WT (n =4), OGT TG (n = 4) and OGA TG (n = 4) animals. Data are represented as mean ± SEM, and significance was determined using a 1 way ANOVA with Tukey's multiple comparison's test (****P<0.0001, ***P<0.001, **P<0.01, *P<0.05, ns=not significant).

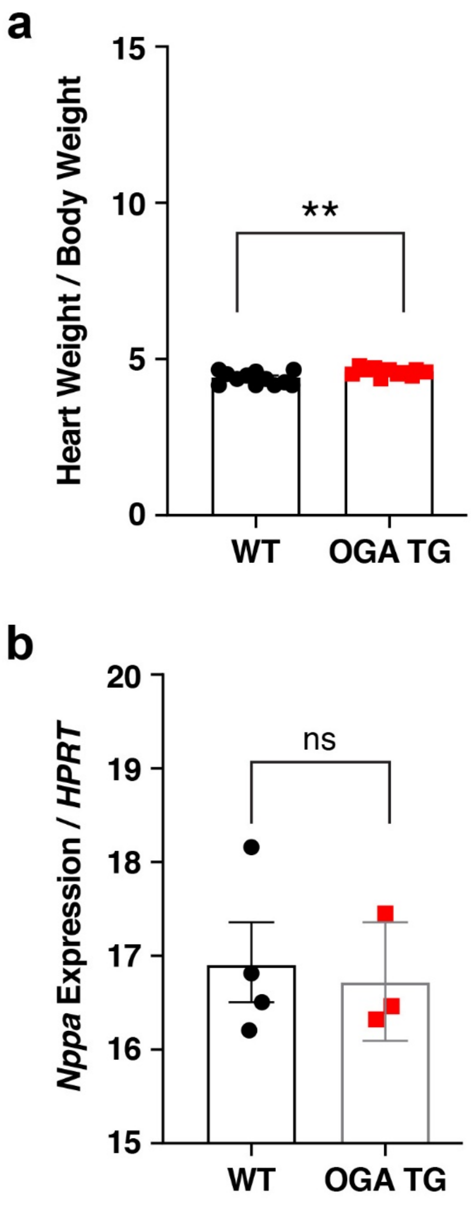


**Fig S2-Supplemental Figure 2**

**OGA TG have increased heart weight but normal expression of heart failure markers**

**a.** Heart weight/body weight for OGA TG compared to WT littermates, WT (n = 12) and OGA TG (n = 10) **b**. Quantification of *Nppa* mRNA expression normalized to Hypoxanthine Peroxidase Reductase Transferase (*Hprt*) in OGA TG (n = 3) and WT (n = 4) hearts at baseline (two independent experiments). Data are represented as mean ± SEM, and significance was determined using a two-tailed Student's *t* test (****P<0.0001, ***P<0.001, **P<0.01, *P<0.05, ns=not significant).

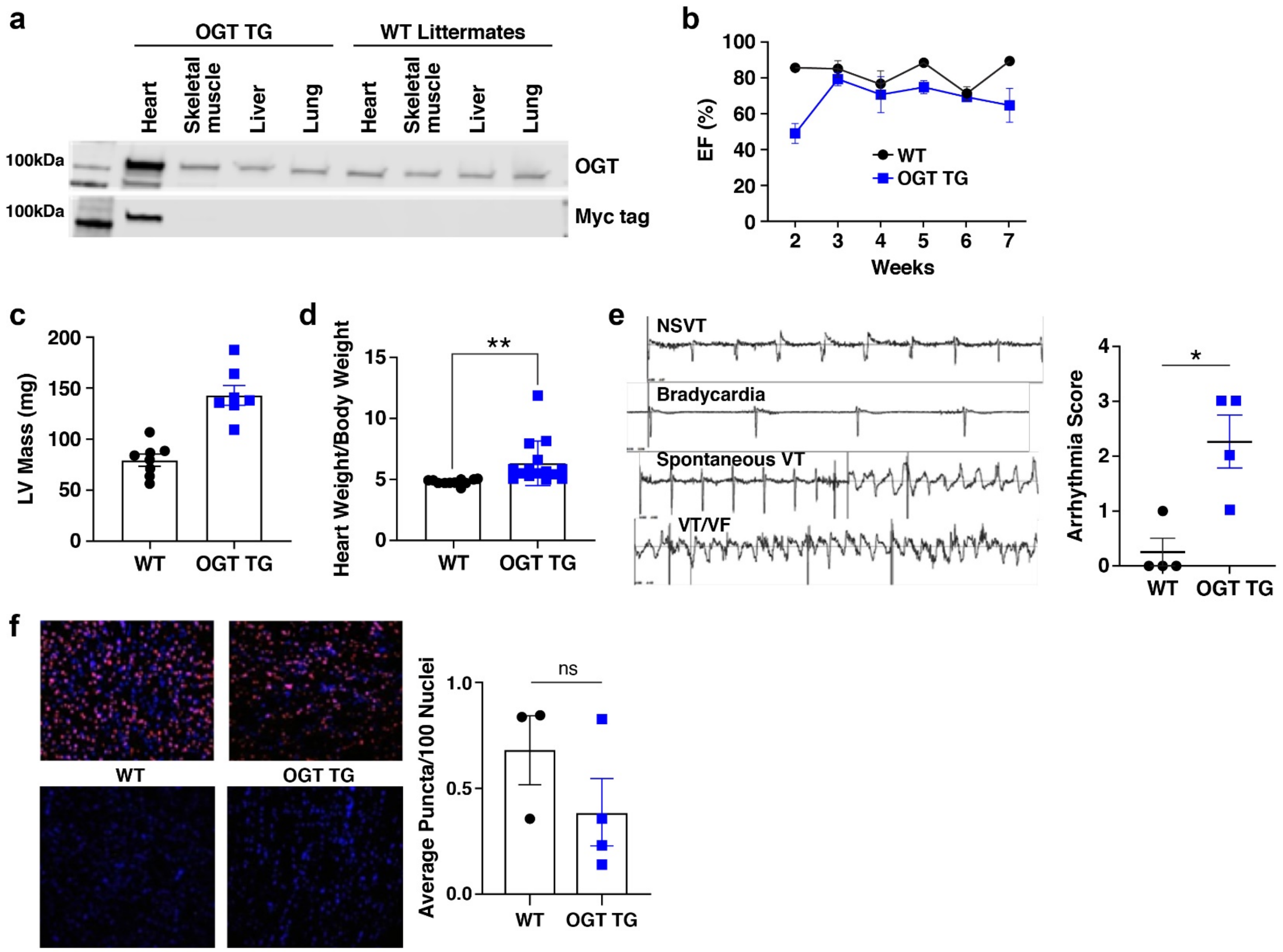


**Fig S3 - Supplemental Figure 3**

**OGT TG mice have dilated cardiomyopathy, and spontaneous arrhythmia in the absence of significant cell death or fibrosis.**

**a.** Western blot for transgene expression in heart, skeletal muscle, liver and lung. Protein isolations from WT and OGT TG mice harvested from tissues of interest. OGT (*O*-GlcNAc transferase). **b.** evaluation of left ventricular function by 2-D echocardiography over 8 weeks in OGT TG (n = 7) and WT (n = 7) **c.** LV (left ventricular) mass calculated from 2-D echocardiography using age and gender-matched mice WT (n= 8), OGT TG (n = 7) **d.** Heart weight/ Body weight for OGT TG (n = 15) versus WT (n = 11) mice. **e.** example electrocardiographic tracings (left) from an OGT TG mouse with a surgically implanted telemeter demonstrating non-sustained ventricular tachycardia (NSVT), bradycardia, sustained ventricular tachycardia (VT) and ventricular fibrillation (VF) and arrhythmia score (right) from WT (n = 4), OGT TG (n = 4) mice at 20 weeks of age **f.** Example TUNEL assay micrographs (left panels) from left ventricular tissue sections obtained from OGT TG and WT hearts and summary data (right panel, WT n=5, OGT TG n =5). Top two images are DNAse treated positive controls in WT (left image) and OGT TG (right image) left ventricular tissue sections. Data are represented as mean ± SEM, significance was determined using a two-tailed t test (****P<0.0001, ***P<0.001, **P<0.01, *P<0.05, ns=not significant).

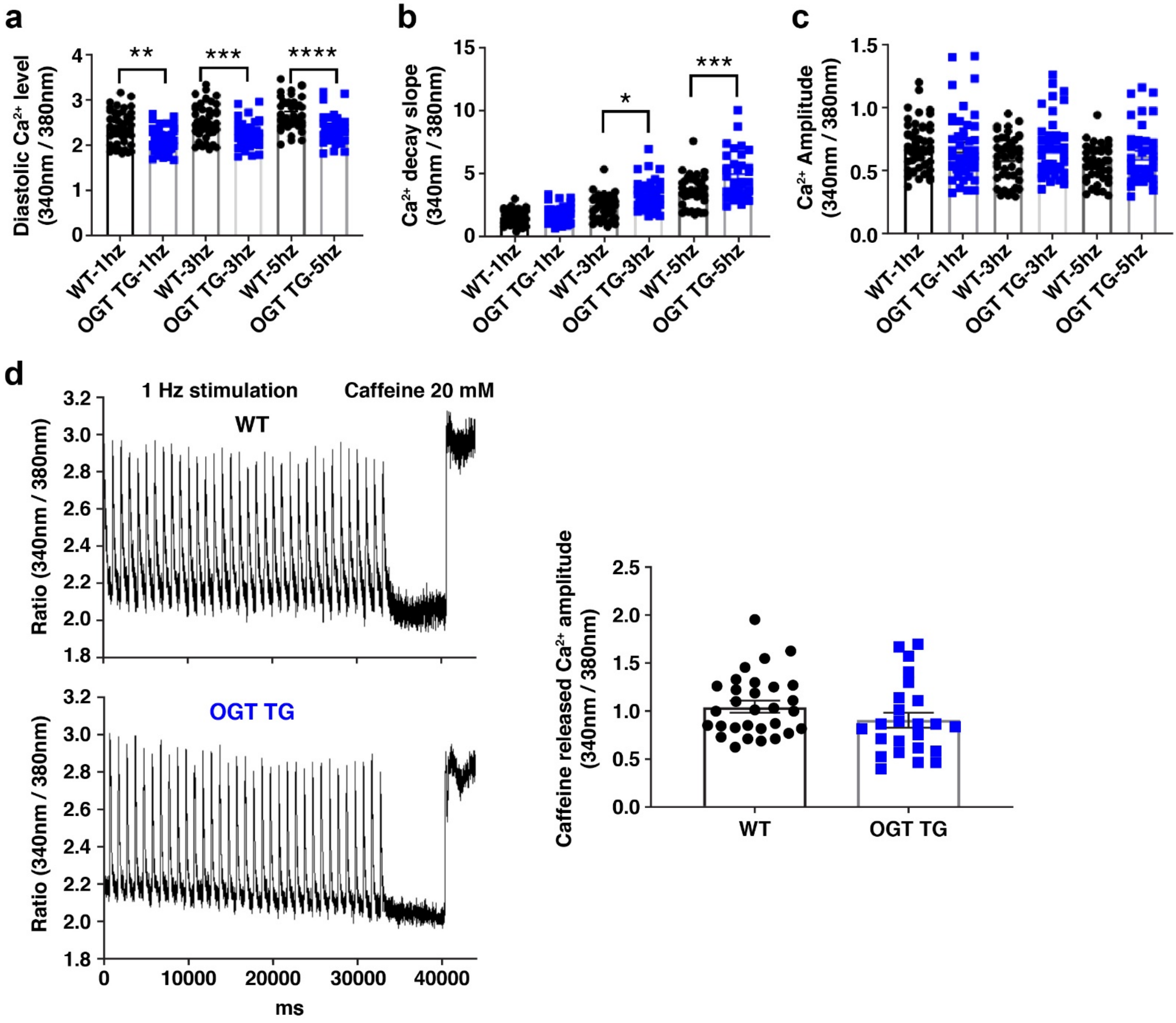


**Fig S4 - Supplemental Figure 4**

**Intracellular $Ca^{2+}$ transients and sarcoplasmic reticulum (SR) $Ca^{2+}$ content in isolated ventricular myocytes**

**a.** Summary data for diastolic $Ca^{2+}$ concentrations in response to 1, 3, and 5 Hz stimulation in ventricular myocytes (n=39-48/group using n=3 mice/group). Date were analyzed with one-way ANOVA and Tukey's multiple comparisons tests. **b**. Summary data for $Ca^{2+}$ transient decay rates and **c**. $Ca^{2+}$ transient amplitudes from cells shown in panel b. Data were analyzed using one-way ANOVA and Tukey's multiple comparisons tests. *p<0.05, ** p<0.01, *** p<0.001, and **** p<0.0001 **d**. Representative recordings of stimulation (1 Hz) induced intracellular $Ca^{2+}$ transients for SR $Ca^{2+}$ loading, and maximal caffeine (20 mM) application induced SR $Ca^{2+}$ release in ventricular myocytes isolated from OGT TG (left lower panel) and WT (left upper panel) mice Summary data for caffeine induced SR $Ca^{2+}$ release (n=24-29 cells/group) isolated from OGT TG (n=2) and WT (n=3) mice.

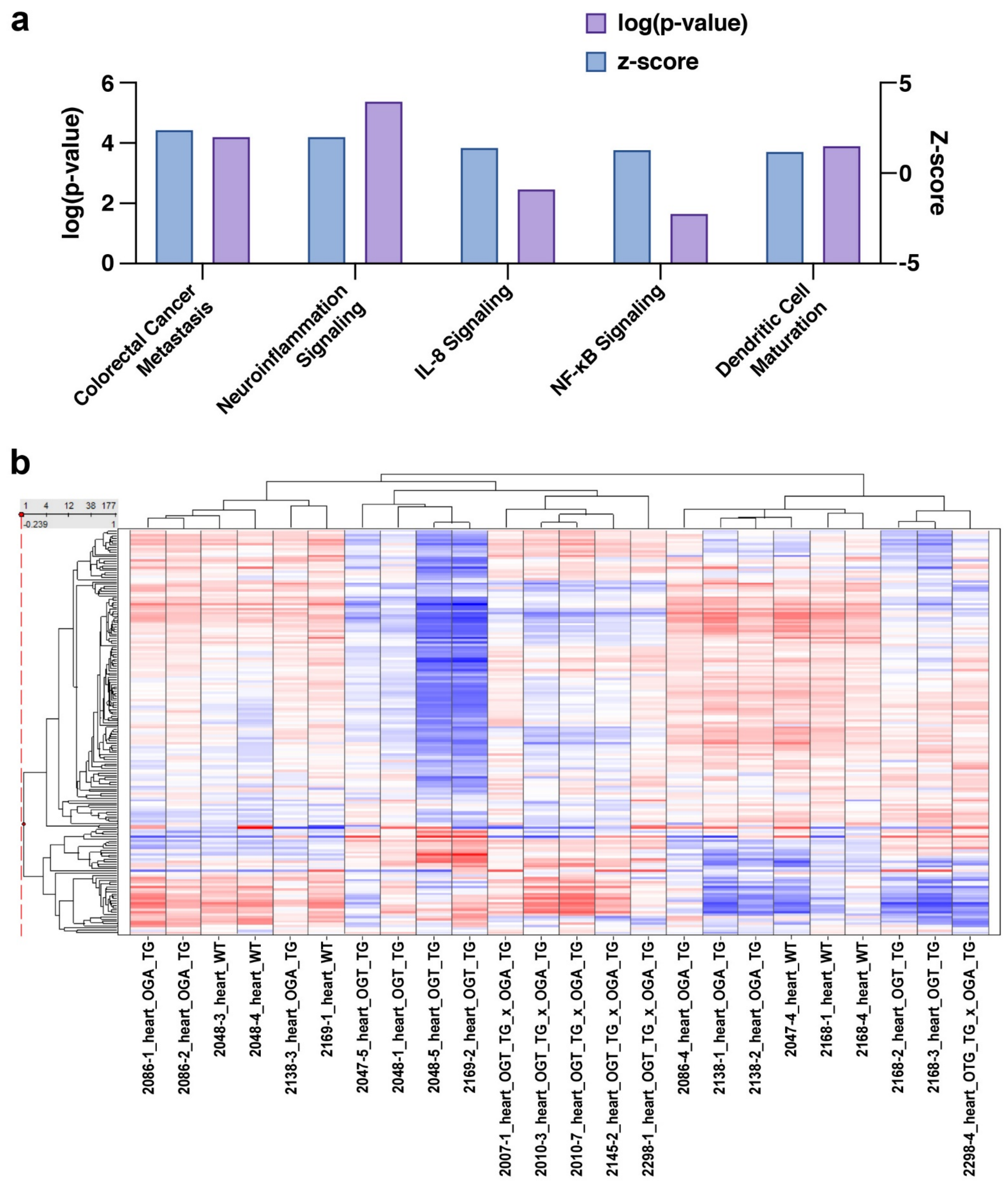


**Fig S5 - Supplemental Figure 5**

**Functional annotation gene clustering and transcriptome landscape between genotypes**

**a**. Functional annotation clusters identified by Ingenuity Pathway Analysis of activated functional pathways between OGT TG and OGT x OGA TG mice. Annotation clusters were identified using default settings with medium stringency (see Methods). **b**. Heat map representing observed mRNA abundance of 177 genes identified in the GSEA hallmark gene set for oxidative phosphorylation, and shown by ANOVA as demonstrating a significant difference between genotypes (n=6 for each genotype). Hierarchical clustering was used to identify coordinated patterns of change.
